# Cryptic genetic structure of the coral host is the primary driver of holobiont assembly in massive *Porites*

**DOI:** 10.1101/2024.01.09.574877

**Authors:** Carly B Scott, Raegen Schott, Mikhail V Matz

**Author notes:** **Corresponding Author:** CB Scott.

## Abstract

The fate of coral reefs in response to climate change depends on their ability to adapt to new environments. The coral animal is buffered from environmental stress by its algal endosymbionts and microbial partners (together, the “holobiont”). However, the flexibility of holobiont community assembly is not well understood, making it difficult to estimate its contribution to coral adaptation. To clarify these processes, we genetically profiled holobiont components (coral, algal symbiont, and microbiome) of massive Porites sampled across two size classes (>2m and <30cm) and ecologically distinct reef sites near Orpheus and Pelorus Islands, Australia. We recovered five major genetic clusters in the coral host, with additional splits within them potentially defining up to ten distinct subclusters. We estimated the relative contributions of the host genetic structure, site, and size class to holobiont community composition. Host genetic structure was the primary driver of both Symbiodiniaceae and microbial communities, indicating strong holobiont specificity in cryptic lineages. In addition, the microbial community was associated with reef site and size class, unlike Symbiodiniaceae that were not significantly affected by either factor. As environmentally segregated, cryptic genetic lineages emerge as a common feature of scleractinian corals, these results emphasize that failure to assess cryptic genetic structure of the coral host may lead to dramatic overestimation of holobiont flexibility.

## Introduction

The rapid rate of climate change poses an immediate threat to coral reefs globally. Environmental change acts on the entire coral holobiont, or the collective unit of the coral animal, its intracellular algal symbionts, and its associated microbial community (Goulet et al., 2020). Studies have posited that specific holobiont communities may increase the resilience of the coral to stress (Voolstra et al., 2021). Thus, corals may keep pace with climate change in one of two ways – through the evolution of the coral host animal and/or through holobiont community reassembly (potentially driven by host evolution). Given the acceleration of global change, acclimatization to future conditions through microbial community shifts has garnered significant attention as a hopeful mechanism for reef persistence (Berkelmans & van Oppen, 2006; Morikawa & Palumbi, 2019; Peixoto et al., 2017).

The underlying mechanisms which regulate coral holobiont assembly are still not fully understood. Without understanding the drivers of coral-associated communities, it is difficult to assess their acclimatization potential. It is likely that genetic, spatial, and temporal factors interact to drive the coral holobiont (Dunphy et al., 2019). Coral taxa exhibit distinct “core” holobiont members in the same environments, implying the functional importance of certain microbial species to the reef (Morrow et al., 2012; Pollock et al., 2018). Similarly, species exhibit geographic variability in their microbial and microalgal associations, indicating a degree of local acclimatization (Hernandez-Agreda et al., 2018).

Over-time studies of the holobiont response to stressors (e.g., bleaching) have yielded conflicting results. Stress events generally induce holobiont reassembly for both Symbiodiniaceae and microbial communities, but the degree and permanence of these shifts is highly variable (Baker, 2003; Glasl et al., 2019; McDevitt-Irwin et al., 2019; Zhu et al., 2022). For Symbiodiniaceae, studies have reported both stable switches to a new symbiont association (Claar, Starko, et al., 2020; Rouzé et al., 2019) and a return to the native state after the stress event (LaJeunesse et al., 2009; Thornhill et al., 2006). The degree of variability of the coral microbiome is also not well established, but the effect of time is often dwarfed by environmental and species-specific pressures (Dunphy et al., 2019; Epstein et al., 2019; Quek et al., 2023).

It has been proposed that coral holobiont dynamics are ungeneralizable and likely geographically and genotypically unique (Dunphy et al., 2019; Hoegh-Guldberg et al., 2007; Stat et al., 2009). While this is possible, most studies fail to integrate long-term (e.g., greater than 5 year) temporal data and individual-level genotyping to disentangle these effects. Thus, the relative roles of host genotype, environment, and time have yet to be resolved at ecologically relevant scales.

Here, we compared host-associated communities massive *Porites* across size classes. The massive *Porites* species complex forms the structure of many Pacific reefs, growing up to 17 meters in diameter and living over 500 years (Brown et al., 2009; Smith et al., 2021). Additionally, massive *Porites* are broadcast spawning species which transmit their Symbiodiniaceae vertically through eggs and generally transmit their microbial community horizontally (Forsman et al., 2020). Thus, we expect the Symbiodiniaceae community to be tightly associated with host genetics and the microbial community to be strongly affected by the local environment.

We sampled small (<30 cm) and giant (>2m and up to 10m) massive *Porites* colonies in 2019 at four sites across Orpheus and Pelorus Islands, Australia. Despite their close proximity, these sites exhibit qualitatively different environments: NE and SE Pelorus are deeper sites with generally high visibility and high wave action, Pioneer Bay is a shallow, nearshore, and highly turbid site, and South Orpheus is also a shallow nearshore site but with fast currents due to its channel location. From this we aimed to, (i) investigate the genetic structure of the coral hosts, and (ii) estimate the relative importance of this genetic structure, site, and size class for the composition of the associated Symbiodiniaceae and microbial communities.

## Methods

### Sampling

*Porites* spp. were sampled around Orpheus and Pelorus Islands, Australia in November 2019. To estimate the effects of size on holobiont structure, we sampled massive adults (>2m diameter) and their smaller counterparts (<30cm diameter; Figure 1). Three samples were taken across each massive individual (one central and two on opposing sides) and one sample was taken from each putative juvenile. Samples were collected from four sites around the island exhibiting different environments (Figure 1). In total, we collected 193 samples from 90 colonies. Coral fragments were fixed immediately after sampling in 100% EtOH and subsequently stored at −80°C.

**Figure 1.**
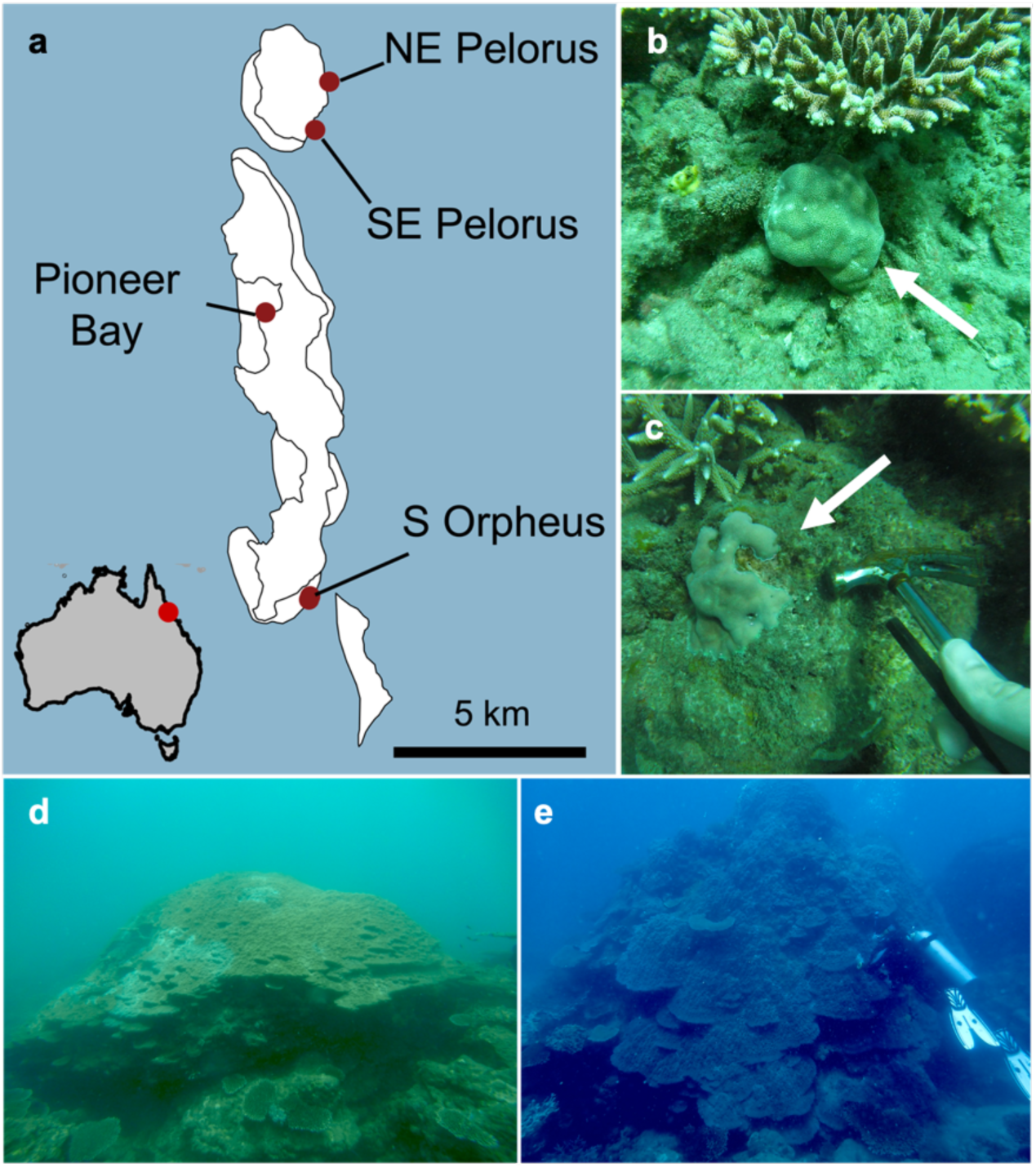
Sampling location and example *Porites* spp. size classes. (a) Map of Orpheus and Pelorus Islands, Australia. These islands are on inner Great Barrier Reef, and their relative location to Australia is marked on the map inset. (b, c) Examples of colonies we considered small *Porites* spp., less than 30 cm in diameter. (d, e) Individuals we considered large *Porites* spp., greater than 2 m in diameter. Individual colonies were not measured due to the large, visible disparity between what we considered size classes.

### DNA Library Preparations

DNA was extracted from all samples using the CTAB method (see supplemental methods). 2bRAD sequencing libraries (https://github.com/z0on/2bRAD_denovo) were constructed for each colony and sequenced using llumina NextSeq 500 SR 75 at the Genomic Sequencing and Analysis Facility at UT Austin. Amplicon sequencing libraries were constructed for all samples. ITS2 amplification used primers from Hume et al., 2015 (SymVarF: CTACACGACGCTCTTCCGATCTGAATTGCAGAACTCCGTGAACC, SymVarR: CAGACGTGT GCTCTTCCGATCTCGGGTTCWCTTGTYTGACTTCATGC); 16S V3/V4 region was amplified using primers from Osman et al., 2020 (16S_341F_Truseq: CTACACGACGCTCTTCCGAT CTCCTACGGGNGGCCTACGGGNGGCWGCAG, 16S_805R_Truseq: CAGACGTGTGCTCT TCCGATCTGACTACHVGGGTATCTAATCC). All amplicon sequencing libraries were amplified with 20-30 PCR cycles, depending on band intensity. Negative controls (MilliQ water) were run with each set of samples and amplification was never observed. Libraries were pooled according to visually assessed gel band brightness. Pooled amplicon libraries were sequenced on the MiSeq PE150.

### Bioinformatic Pipeline

Reads were trimmed of adaptors and deduplicated using custom 2b-RAD scripts (https://github.com/z0on/2bRAD_denovo). Reads containing base calls with quality less than Q15 were removed using *cutadapt* (Martin, 2011). Cleaned reads were mapped to a combined reference of the *Porites lutea* genome (Robbins, 2019), *Symbiodinium* sp. (Shoguchi et al., 2018), *Breviolum minutum* (Shoguchi et al., 2013), *Cladocopium* sp. (Shoguchi et al., 2018), and *Durusdinium* sp. (Dougan et al., 2022) genomes using *bowtie2* with default local settings (Langmead & Salzberg, 2012). Mapped reads were filtered for a mapping quality >30 and genotyping rate >50%, and only reads which mapped to *Porites lutea* were retained. Between-sample genetic distances were calculated as identity-by-state (IBS) based on single-read resampling in ANGSD (Korneliussen et al., 2014).

At this point we subset the data to remove technical replicates, colony replicates, and clones based on hierarchical clustering, retaining one individual per genotype (N=65). After genetically identical samples were removed, the genetic distances were calculated again in ANGSD with the same filtering scheme. At this point, we used ‘pcangsd’ to calculate the most likely number of admixture clusters in our host samples (Meisner & Albrechtsen, 2018).

### Host Genetic Analysis

Initial analysis revealed highly distinct genetic groups within *Porites* spp, even within admixture clusters identified by PCAngsd. To estimate the most likely number of genetic subclusters, we implemented a Bayesian model-based clustering algorithm from the ‘mclust’ R package on our ANGSD-generated IBS matrix (Scrucca et al., 2016). We chose to use the number of clusters (N=10) with the highest Bayesian Information Criterion in subsequent analysis. This visually corresponded well with the hierarchical clustering tree generated from the genetic distance matrix and was concordant with the PCAngsd results (mclust-identified subclusters were always contained within larger admixture groups). Finally, we calculated pairwise *F_ST_* between each genetic subcluster using realSFS.

We used RDA/PERMANOVA analysis to determine the relative contribution of site and size class to *Porites* spp. genetic structure. Using the ‘adonis2’ function from the R package ‘vegan’ (Oskansen et al., 2022) we tested for significance of age and site in predicting genetic distance between individuals using ‘method = marginal’ (genetic distances ~ Site + Size Class + Site*Size Class).

We were additionally interested in patterns which may arise *within* admixture groups and conducted a separate analysis on each admixture group. This analysis started at the ANGSD stage, where a new IBS matrix was generated for the members of each admixture group. We then determined the role of site and size class in structuring the within-group genetic variation through PERMANOVA analysis, using ‘adonis2’ with ‘method=marginal’, with model: Genetic Distance ~ Site + Size Class.

### Symbiont and Microbe ASV Prefiltering

We used the same basic computational workflow for both 16S and ITS2 sequences. Primers and adapters were trimmed from reads with cutadapt. For microbes, likely host contamination was removed from our samples using METAXA2 (Bengtsson-Palme et al., 2015). We only retained reads classified as bacterial or archaeal in origin.

### Determination of Symbiont ASVs

Symbiodiniaceae ITS2 locus are difficult to deal with due to their intragenomic variation and multicopy nature. While the SymPortal (Hume et al., 2019) framework has been proposed to handle these challenges, our reads did not have sufficient overlap between the left and right reads of the pair to be retained through SymPortal prefiltering steps. Thus, ITS2 sequence variants were identified via the DADA2 pipeline in R statistical software (Callahan et al., 2016). To deal with the high degree of intragenomic variation expected in Symbiodiniaceae ITS2 sequences, we calculated the correlation of ASV presence/absence across samples. Under the logic that highly correlated ASV sequences likely correspond to within-species intergenomic variation, rather than additional species diversity, we clustered any ASVs with correlated presences higher than 0.8 into a single “ASV group” which was used in downstream analyses.

To assign taxonomy to the ITS2 ASVs, we constructed a custom blast database from the SymPortal DIV database (updated on 2024-02-13). We then blasted our ASV sequences against this database, keeping the best 30 hits per ASV. From the blast classifications, we retained only the subtype of each strain’s entry in the SymPortal database (e.g., C15 rather than C15au). We filtered the blast results for hits with an e-value < 1e-100 and >95% match to the target sequence. At this threshold, some queries still had multiple best hits. For these queries, we determined if there was consensus in their best hits at the subtype-level (e.g., all C15, but different strain identifiers). We assigned a query ASV this subtype if there was over 90% consensus in the blast hits with e-value < 1e-100 and >95% match to the target. For ITS2 ASVs not meeting this criterion, we did not assign taxonomy.

To aggregate our colony-level replicates, we calculated the sum of ASV groups across replicates to gain a “colony-averaged” community. ASV groups were retained if they were present in at least three colonies and had >100 reads across samples. Similarly, colonies were retained if they had >3 nonzero ASVs present and > 1000 total sequenced reads.

### Determination of Microbial ASVs

The DADA2 pipeline was also used on trimmed microbial reads for denoising and ASV classification. We assigned taxa to ASVs in DADA2 using the ‘assignTaxa’ function, using the Silva version 138.1 SSU reference database (DADA2 formatted database taken from: https://zenodo.org/records/4587955; Quast et al., 2013).

Similar to the ITS2 ASV filtering, we obtained a colony-level community measure by summing across ASVs in colony replicates. We only retained ASVs present in > 3 colonies with a minimum of 100 reads across colonies. We only kept colonies with > 1000 total sequenced reads and >3 nonzero ASVs present.

### Community-level Analysis

The same analytical pipeline was used for both ITS2 and 16S ASVs in parallel. Given we wanted to evaluate the role of host genetics in structuring the holobiont community, we only retained colonies which had corresponding 2bRAD sequencing data. ASV abundances were then standardized across colonies using the Hellinger method using the ‘decostand’ function from vegan. Bray-Curtis distances between colonies were calculated from Hellinger-standardized abundances using ‘vegdist’.

We evaluated the role of host genetic structure, site, and size class in two different ways. First, we conducted a PERMANOVA analysis using vegan’s ‘adonis2’ function, with option “method=marginal”. We tested two models for each community distance matrix: Holobiont Community Distance ~ Host Genetic Subcluster + Site + Size Class, and, Holobiont Community Distance ~ Host admixture group + Site + Size Class. In parallel, we conducted an RDA-forest analysis (Black et al., 2022; https://github.com/z0on/RDA-forest). RDA-forest is an application of gradient forest (a multidimensional extension of random forest; Ellis, Smith, & Pitcher, 2012) to multi-dimensional data, with the goal of predicting PcoA structure. We first constructed an unconstrained PCoA from community distances using vegan’s ‘capscale’ command. The first fifteen principal coordinates were used as the response matrix, and the predictor matrix included host admixture group (or subcluster), site, and size class. Categorical predictors with multiple levels (site and genetic cluster) were converted into dummy quantitative variables with values 1 and 0, with one variable per predictor level. Each random forest model used 1500 bootstrapped trees and max variable permutation level of log2(N_Individuals_* 0.368/2). Then, for multi-level categorical predictors, we summed the weighted importance across dummy variables representing their levels to obtain overall importance. Similar to our PERMANOVA analysis, we ran two versions of this model, one with admixture group as a predictor and one with genetic subcluster as a predictor.

To determine what structured communities *within* admixture groups, we ran separate PERMANOVA analyses for each group, with ‘method=marginal’ and model: Community Distance ~ Genetic Subcluster + Site + Size. When there were no genetic subclusters within an admixture group, this predictor was omitted. We set our significance threshold at *p* < 0.01, correcting for multiple hypothesis testing.

### 16S Differential Abundance Analysis

Differential abundance of microbes was calculated using DESeq2 on the raw abundance matrix with the experimental design, abundance~Genetic Subcluster + Site + Age (Love, Huber, & Anders, 2014). We designated α < 0.01 as the cutoff to determine which taxa were significantly differentially abundant between comparisons.

All results were visualized in R with ggplot2. Scripts for replicating this analysis can be found at https://github.com/cb-scott/PoritesHolobiont_Final and archived on Zenodo (doi:10.5281/zenodo.10384279).

## Results

### Porites spp. host genetic structure predicted by site

Initial population genetic analysis suggested a high degree of clustering/sub clustering. We identified five admixture groups within the individuals we sampled (Figure 2a,b; Figure 3a). However, Bayesian model-based clustering returned the optimal number of clusters as ten, corresponding well to the hierarchical clustering tree generated between samples (Supplemental Figure 1–2). These groups represent “subclusters” of the identified admixture clusters (Figure 3b-f). Calculation of weighted pairwise *F*_ST_ further supported genetic distinction between these ten groups (minimum pairwise *F*_ST_: 0.14, maximum: 0.55; Supplemental Figure 3).

**Figure 2.**
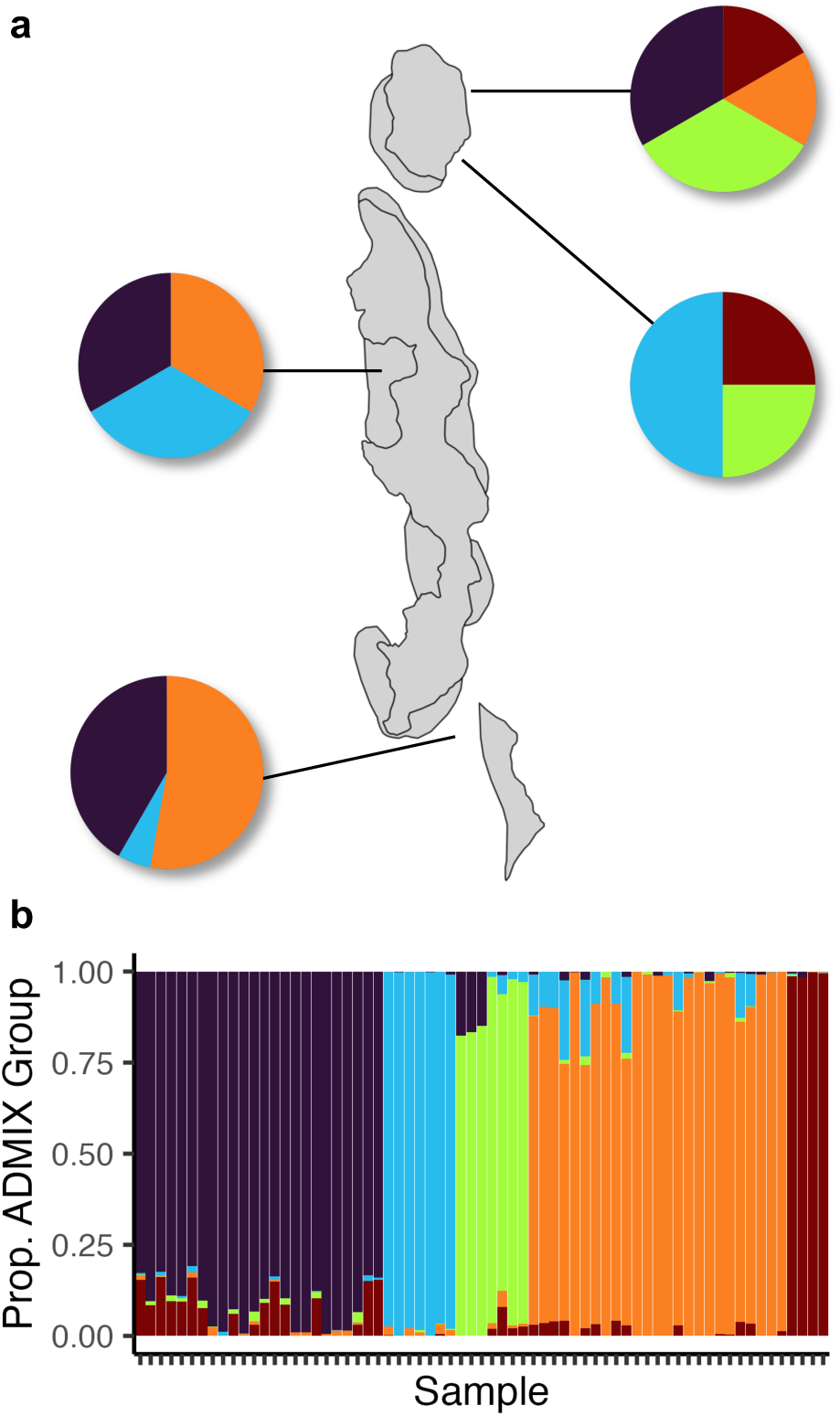
Host genetics structured by site. (a) Proportion individuals from each admixture group at each site. (b) admixture plot representing all colonies sequenced, with each vertical bar representing one sample. Ordered by maximum group membership.

**Figure 3.**
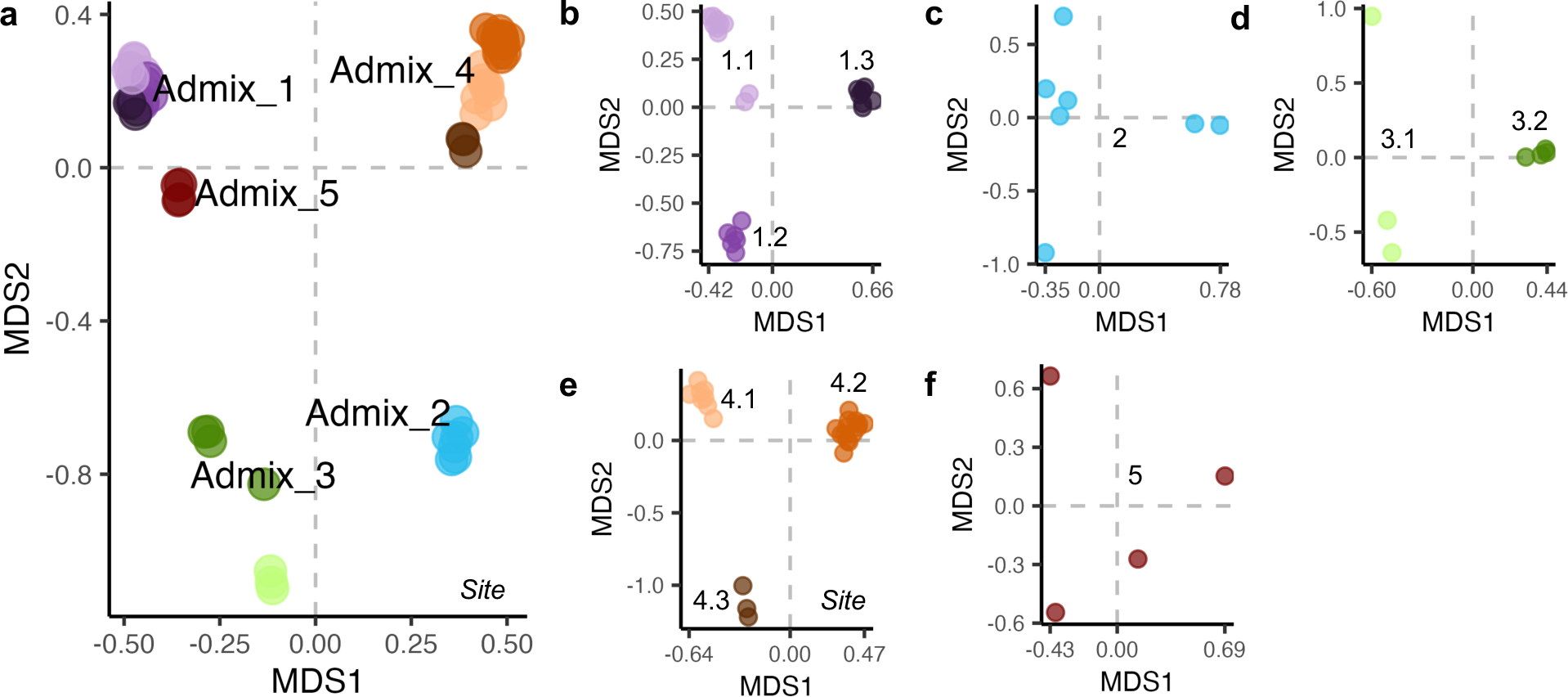
A large degree of cryptic genetic structure exists with *Porites spp* admixture groups. (a) PCoA of genetic distances colored by assigned admixture group (main color) and Bayesian-assigned subcluster (shades of main color). (b-f) Genetic substructure of each admixture group with subclusters indicated by labels on plot. Subcluster numbering corresponds to admixture group. Variables names on plots indicated significant predictors of genetic structure by PERMANOVA

Through a PERMANOVA analysis, we determined that site was the only significant (at the *p < 0.05* level) factor in determining the overall genetic structure of Orpheus and Pelorus Island *Porites* spp. (Figure 2a, Figure 3a, Table 1). We additionally ran this analysis on each admixture group separately. Only admixture group 4 was significantly structured by site (*p < 0.01*), when our threshold p-value was corrected for multiple hypothesis testing (Supplemental Table 1).

**Table 1.**
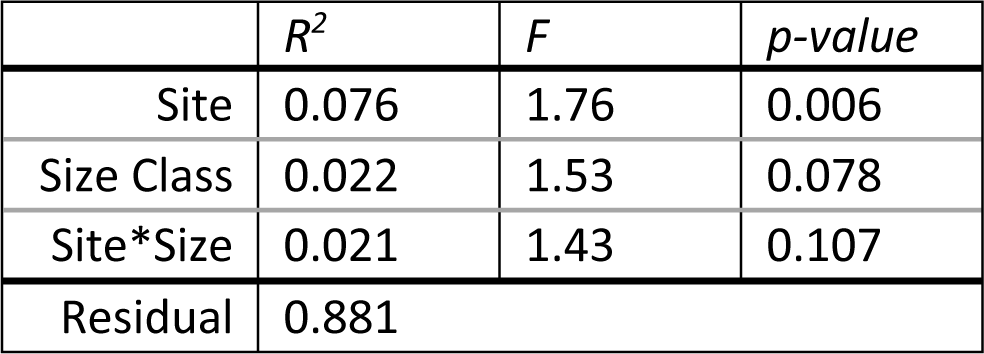
Site structures *Porites* spp. host genetics. PERMANOVA results for the model, Genetic Distance ~ Site + Size Class + Site*Size Class, with the marginal importance of each variable given.

| | $R^2$ | $F$ | $p$-value |
| --- | --- | --- | --- |
| Site | 0.076 | 1.76 | 0.006 |
| Size Class | 0.022 | 1.53 | 0.078 |
| Site*Size | 0.021 | 1.43 | 0.107 |
| Residual | 0.881 |  |  |

### Holobiont community composition

The vast majority of the Symbiodiniaceae ASVs we identified as C15 (Figure 4). For large *Porites* spp., the Symbiodiniaceae community from colony-level replicates had significantly lower variance than within-admixture or within-site groups (Supplemental Figure 4). Microbial communities were dominated by ASVs assigned to Alphaproteobacteria and Gammaproteobacteria (Figure 5). Similar to the Symbiodiniaceae community, within colony variance was significantly lower than within-site or within-admixture variance (Supplemental Figure 5).

**Figure 4.**
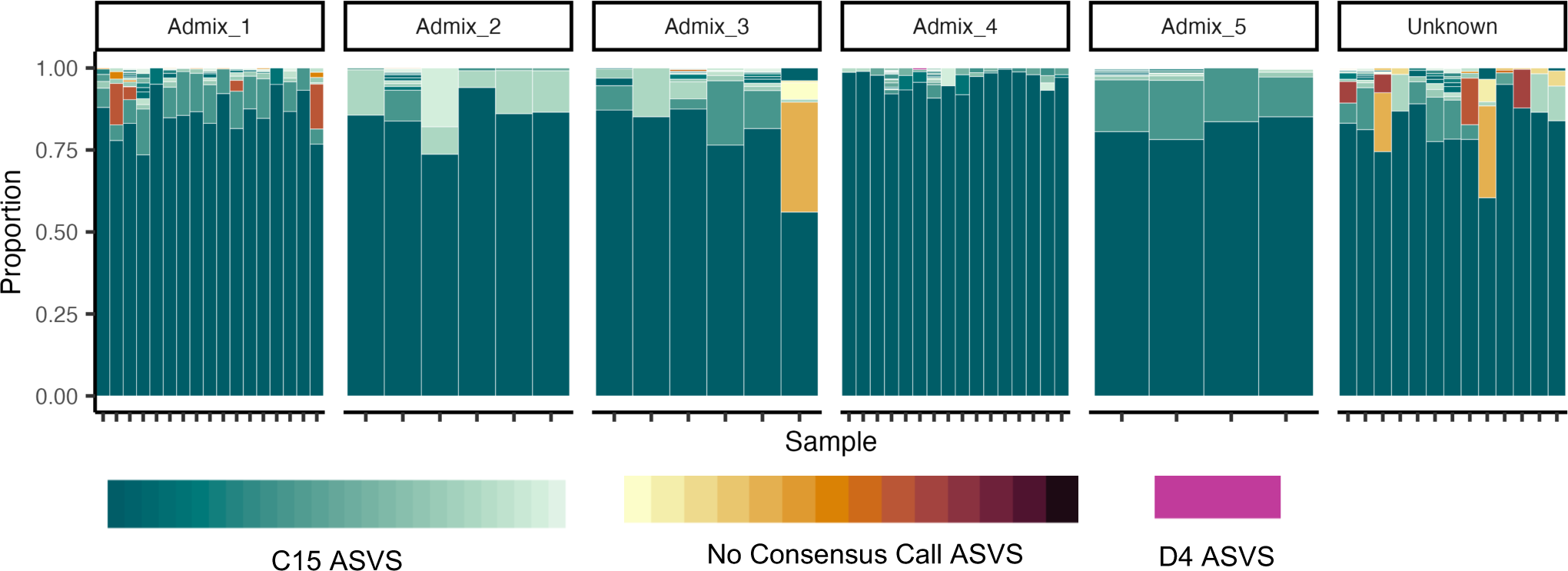
C15 ASVs dominate Symbiodiniaceae community. Stacked bar plot where bars represent coral colonies and colors show assigned taxonomy for each “ASV group”. Blue colored bars are ASV groups identified as a subtype of C15 Symbiodiniaceae. Orange bars show ASV groups which had no consensus call based on our custom blast. Samples are organized by the colony’s assigned admixture group (panels). Those in the “unknown” group did not have sufficient host genetic data to carry into downstream analysis.

**Figure 5.**
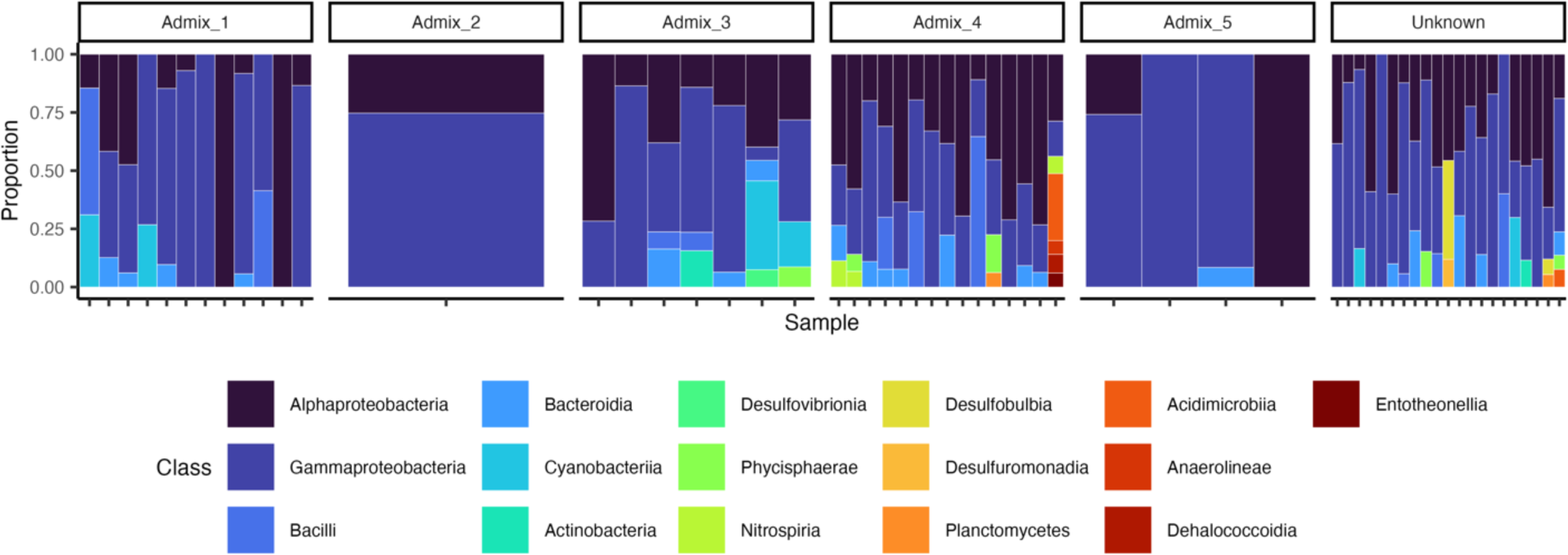
Porites microbial community composition across admixture groups. Stacked bar plot where bars represent coral colonies and colors show assigned taxonomy at the class-level. Only taxa making up at least 5%, but no more than 90%, of the microbial community are shown. Samples are organized by the colony’s assigned admixture group (panels). Those in the “unknown” group did not have sufficient host genetic data to carry into downstream analysis.

### Symbiodiniaceae community more structured by host genetics than microbial community

We compared the relative importance of host genetics, size class, and site in structuring the Symbiodiniaceae and microbial community using an RDA-forest model. For both holobiont members, the host’s genetic subcluster primarily drove community variation, followed by site, then size class (Figure 3c, Supplemental Tables 2 – 3).

RDA-forest supported a stronger link between the host’s cryptic genetic subcluster and Symbiodiniaceae community than the microbial community. For the Symbiodiniaceae, of the total gradient forest model importance (total importance = 0.155), 81% of the importance was generated by genetic subcluster (weighted variable importance = 0.155), 11% was generated by site (weighted variable importance = 0.021), and 8% was generated by size class (weighted variable importance = 0.016; Supplemental Table 2). At the same time, for the microbial community, of the total gradient forest model importance (total importance = 0.425), only 50% of the importance was generated by genetic subcluster (weighted variable importance = 0.212), 35% was generated by site (weighted variable importance = 0.148), and 11% was generated by size class (weighted variable importance = 0.050; Supplemental Table 3). These results were additionally supported by a separate PERMANOVA analyses (Tables 2 – 3) and. All results were qualitatively the same when admixture group was used as a predictor rather than genetic subcluster (Supplemental Tables 4 – 7).

**Table 2.** PERMANOVA results for drivers of Symbiodiniaceae community structure support genetic subcluster as most important predictor. Host genetics have the strongest (and only significant) effect on symbiont community, followed by sampling site, then size class.

| Predictor | $R^2$ | $F$ | $p$ |
| --- | --- | --- | --- |
| Subcluster | 0.527 | 4.78 | 0.001 |
| Site | 0.034 | .936 | .512 |
| Size Class | 0.012 | 0.98 | .365 |
| Residual | 0.428 |  |  |

**Table 3.**
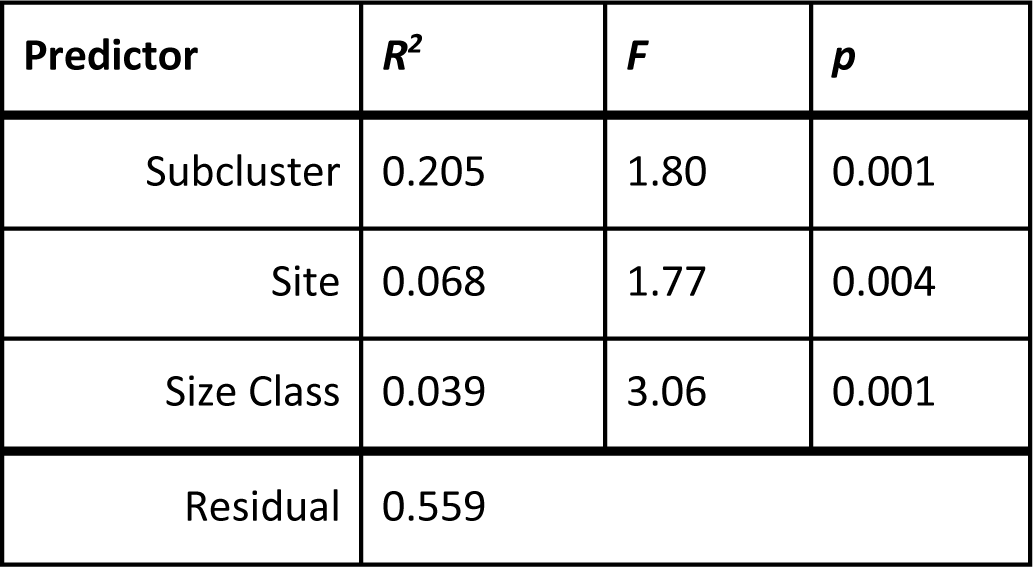
PERMANOVA results for drivers of microbial community structure show a significant role of genetics, site, and age. Host genetics have the strongest effect on the microbial community, followed by sampling site, then age.

### Differentially abundant microbial taxa between groups

We found find 29 significantly differentially abundant bacterial taxa between size classes and/or between sites (Supplemental Figure 6 – 7). Most notably, some strains of *Vibrio* spp. and *Endozoicomonas* spp. were differentially abundant between sites. Additionally, some strains of *Vibrio* spp. were differentially abundant between size classes.

## Discussion

We found a high degree of cryptic genetic structure in massive Porites. This is unsurprising, as previous studies have revealed complex genetic substructure within the species complex (Forsman et al., 2020; Starko et al., 2023). We identified 10 distinct genetic groups in this study from just two neighboring islands in the Indo Pacific. Likely, the distinct admixture groups recovered reflect the sampling of multiple species of *Porites* spp., as they are notoriously difficult to identify in the field. Still, since there are only three named species of massive *Porites*, some of our 10 genetic lineages likely reflect additional cryptic genetic diversity within the nominal species. Much like previous studies, these cryptic lineages are associated with specific geographical locations, hinting at an evolutionary mechanism linked to diverse environmental conditions.

For both the Symbiodiniaceae and microbial communities, host genetic subcluster was the most important factor in determining community structure (Figure 6c). This aligns with recent work showing that cryptic species of *Porites lutea* host distinct holobionts (Grupstra et al., 2024). This was most stark for the vertically transmitted Symbiodiniaceae community, where over 80% of the predictive power in our RDA-forest model came from genetic subcluster. Combined with the PERMANOVA results, we did not find support for site-specific or size-specific Symbiodiniaceae communities. This advances evidence for long-term coevolution between *Porites* spp. and their symbiotic algae (Hoadley et al., 2021; Starko et al., 2023). All coral colonies were dominated by symbiont strain C15, so differences between genetic subclusters are being driven by background strains (Figure 4). It is possible that hosting different background Symbiodiniaceae communities may advantage the host during stress-induced “symbiont shuffling” (e.g., Baker 2003; Coffroth et al., 2022; Starko et al., 2023). However, given host genetic structure is associated with site (Figure 2a, Table 1), this pattern may simply reflect the environmental abundance of Symbiodiniaceae.

**Figure 6.**
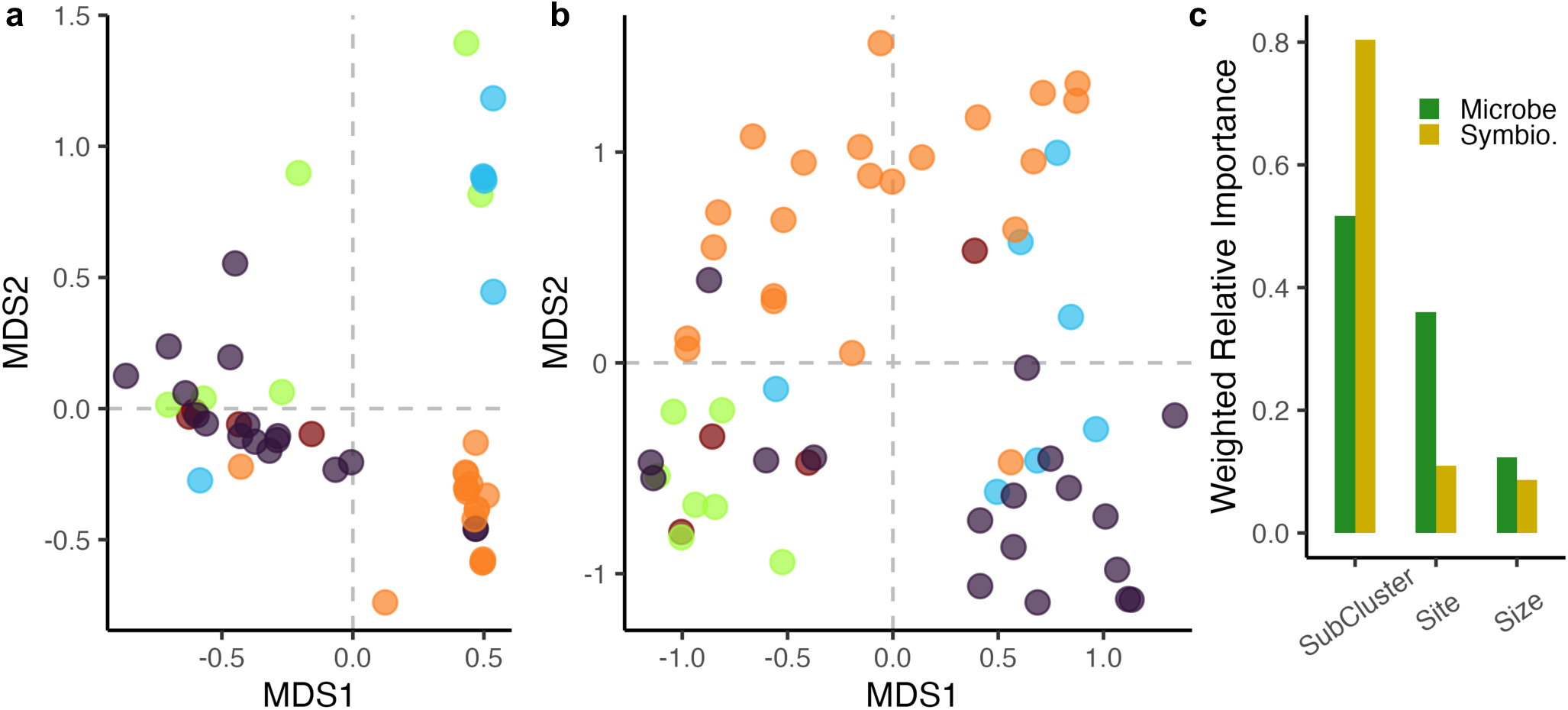
Host genetics is the primary driver of Symbiodiniaceae and microbial community composition. (a) PCoA of Symbiodiniaceae community structure, based on Bray-Curtis distances, colored by host colony admixture group. (b) PcoA of microbial community structure, based on Bray-Curtis distances, colored by host colony admixture group. (c) RDA-forest model results giving weighted relative importance for Symbiodiniaceae (gold) and microbial (green) communities. In brief, weighted relative importance represents the proportion of variance explained by a given variable out of the total variance explained by the model. For both microbes and Symbiodiniaceae, genetic subcluster is the most important predictor of community structure. We find qualitatively similar results using admixture group as a predictor in the RDA-forest model (Supplemental Tables 4 – 7) and through PERMANOVA analysis (Tables 2 – 3).

Previous work has identified site-specific, age-dependent microbial diversity patterns in *Porites lutea* (Wainwright et al., 2020). We build on this work, finding a significant effect of site and size class in determining the microbial community structure of massive Porites (Figure 6c, Table 3). However, we note that while Wainwright et al. used corallite structure to assign all samples to *P. lutea*, our results suggest there is likely additional genetic substructure within the named Porites species. We find that genetic subclusters can be structured by site (Figure 3e), thus these site-specific patterns may partially reflect genetic associations with specific microbial communities. Likely, as we show here, site-specific factors are trumped by cryptic genetic lineage (Figure 6c).

Similarly, the size class differences we found in the microbiome are small, but significant, compared to the role of host genetics. Size class specific communities may reflect changes in the microbial community with host age, and/or morphology-driven changes. These differences may be driven by neutral (to the host) ecological processes, such as priority effects (McIlroy et al., 2019); host selection for a specific microbiome over time (Pollock et al., 2018); or colony geometry (Morrow et al., 2022). Evaluating the roles of these drivers should be done controlling for genetic lineage, as fine-scale differences may be obscured by lineage-specific communities.

We found some microbial community differences to be driven by known functionally important taxa (Supplemental Figures 6 - 7) One such case is the differential abundance of *Vibrio* species between large and small colonies. *Vibrio spp*. is an opportunistic pathogen (Munn, 2015) and has been associated with coral bleaching (Ben-Haim et al., 2003). This suggests that particular *Vibrio* strains may be age-class specific pathogens, or, more likely to invade age-specific communities. Another case is between-site differences in *Endozoicomonas* spp. abundance. *Endozoicomonas* has been suggested to be a beneficial microbial symbiont in corals which can modulate environmental stress (Peixoto et al., 2017; Tandon et al., 2022). These site-specific differences may support a functional role of *Endozoicomonas* in certain environments.

Above all, our results stress that failure to assess cryptic genetic structure may lead to the overestimation of holobiont flexibility. Cryptic species complexes are emerging as a common feature of scleractinian taxa and often segregate by site/environment (Fifer et al., 2022; Forsman et al., 2020; Ladner & Palumbi, 2012; Rippe et al., 2021). Thus, site-specific patterns in the holobiont may not reflect flexible environmental associations, but rather genetically determined, lineage-specific communities. It is imperative to genotype hosts in holobiont studies, as morphologically similar – but genetically divergent – sympatric corals are likely associated with unique microbial/microalgal communities.

## Conclusion

We demonstrated cryptic genetic structure is the most important factor in determining both coral-associated Symbiodiniaceae and microbial communities. Local environment and size of the host have much less effect, although the microbial community is more responsive to local environment than Symbiodiniaceae. This strong fidelity to host genetic clusters suggests the holobiont components cannot easily evolve independently from the host. Thus, underlying host genetic structure must always be weighed in determining the acclimatization potential of the holobiont to future conditions.

## Acknowledgements

This research was supported by NSF grant DGE-2137420 to C.B.S, NSF grant IOS-1755277 to M. V. M., funding from the University of Texas Department of Integrative Biology to C.B.S, and funding from the International Women’s Fishing Association to C.B.S. The data analysis has been performed using facilities of the Texas Advanced Computing Center (TACC). We thank Kristina Black, Greg Torda, and JP Rippe for their help collecting these samples and their camaraderie in the field, and, the staff of Orpheus Island Research Station for supporting the logistics of this project.

## Data Availability

All sequencing (2bRAD and 16S/1TS2 amplicon) data generated can be found at NCBI BioProject PRJNAXXX. All scripts and intermediate products (including metadata) used to complete analysis can be found at https://github.com/cb-scott/PoritesHolobiont_Final or archived at doi: XXX.

## Author Contributions

Conceptualization: CBS and MVM; Data curation: CBS and RS; Formal analysis: CBS and RS; Funding acquisition: CBS and MVM; Investigation: CBS and RS; Methodology: MVM; Project administration: MVM; Resources: MVM; Software: all; Supervision: MVM; Validation: MVM; Visualization: CBS; Writing – original draft: CBS; Writing - review & editing: CBS and MVM.

## Supplemental Figures

**Supplemental Figure 1.**
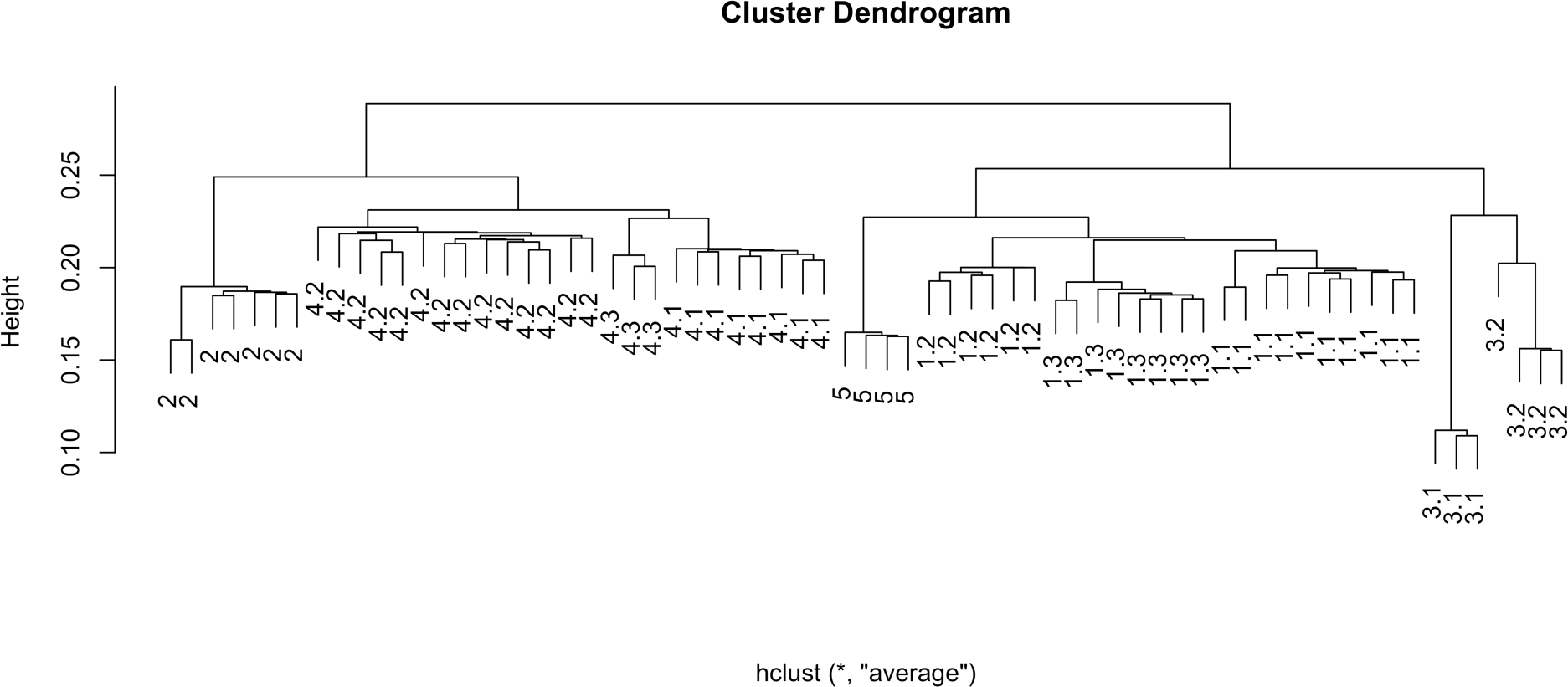
Hierarchical clustering results of genetic data. Hierarchical clustering of ANGSD identity by state matrix aligns well with Bayesian-assigned genetic clusters (leaf labels).

**Supplemental Figure 2.**
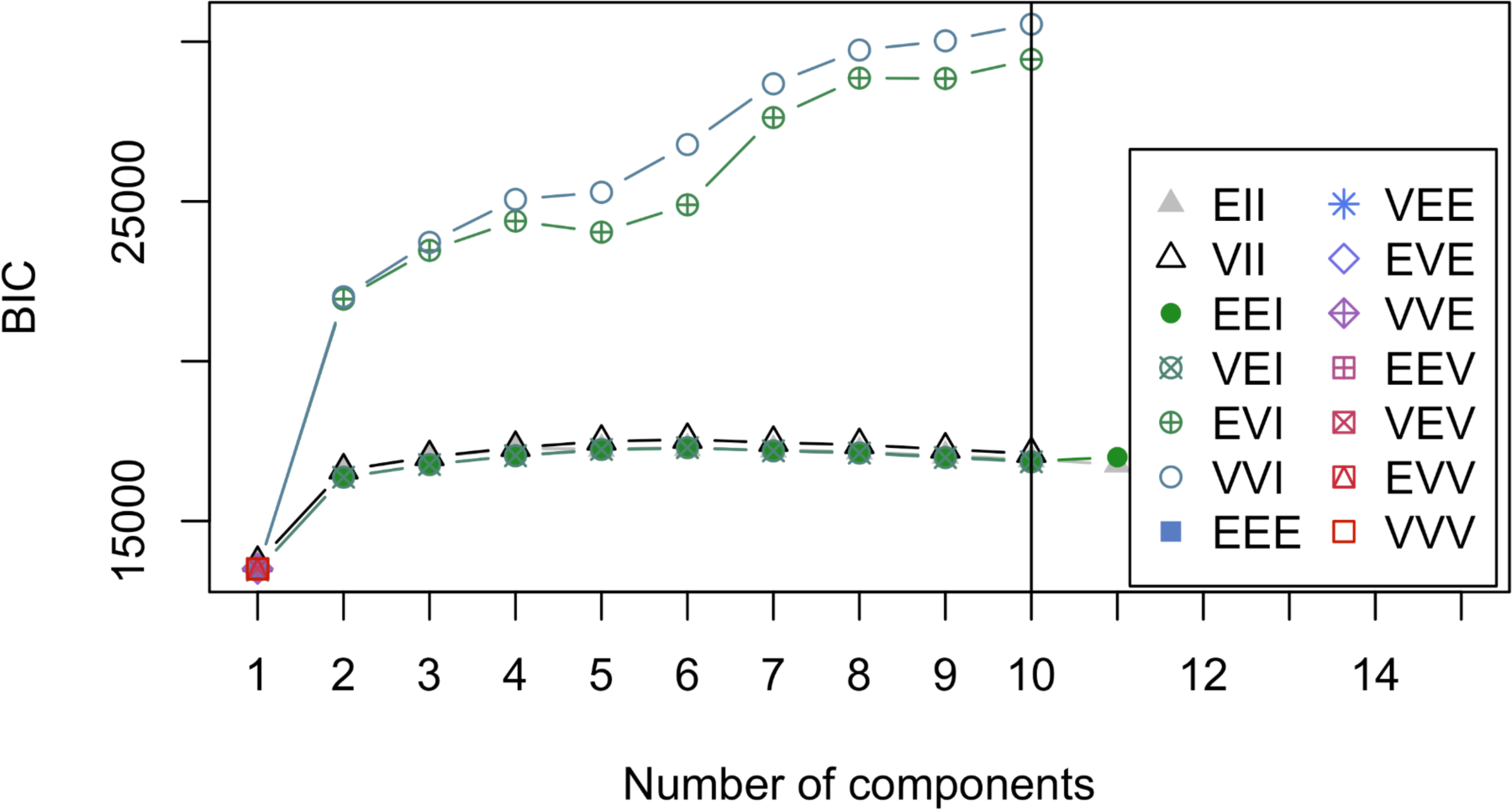
Bayesian Information Criterion used for choosing optimum number of clusters. BIC is maximized at 10 clusters, the value chosen for subsequent analyses.

**Supplemental Figure 3.**
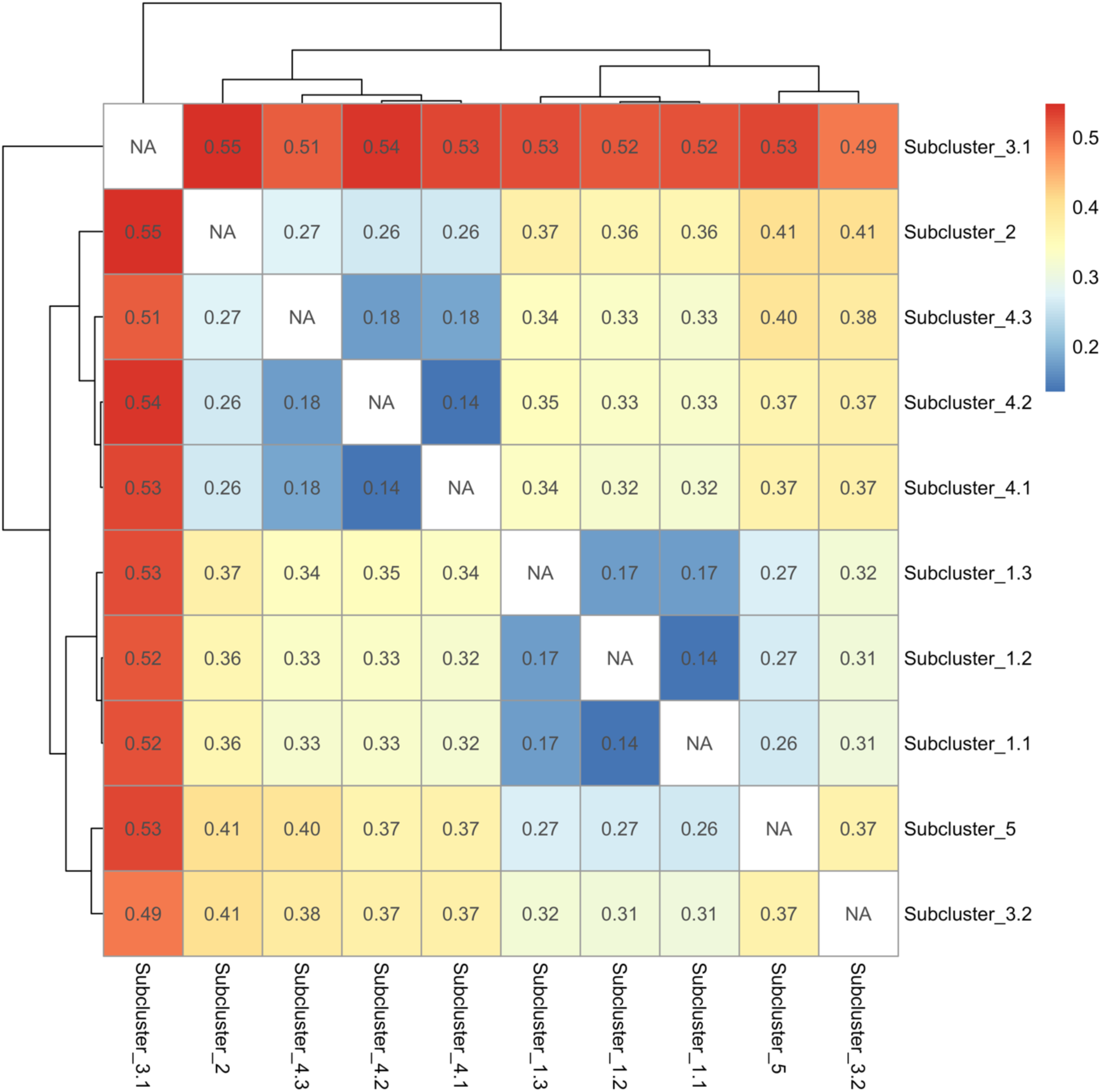
Pairwise *F_ST_* between Bayesian-identified genetic clusters. Calculations of pairwise F_ST_ between clusters indicate a high degree of genetic divergence between cryptic lineages.

**Supplemental Figure 4.**
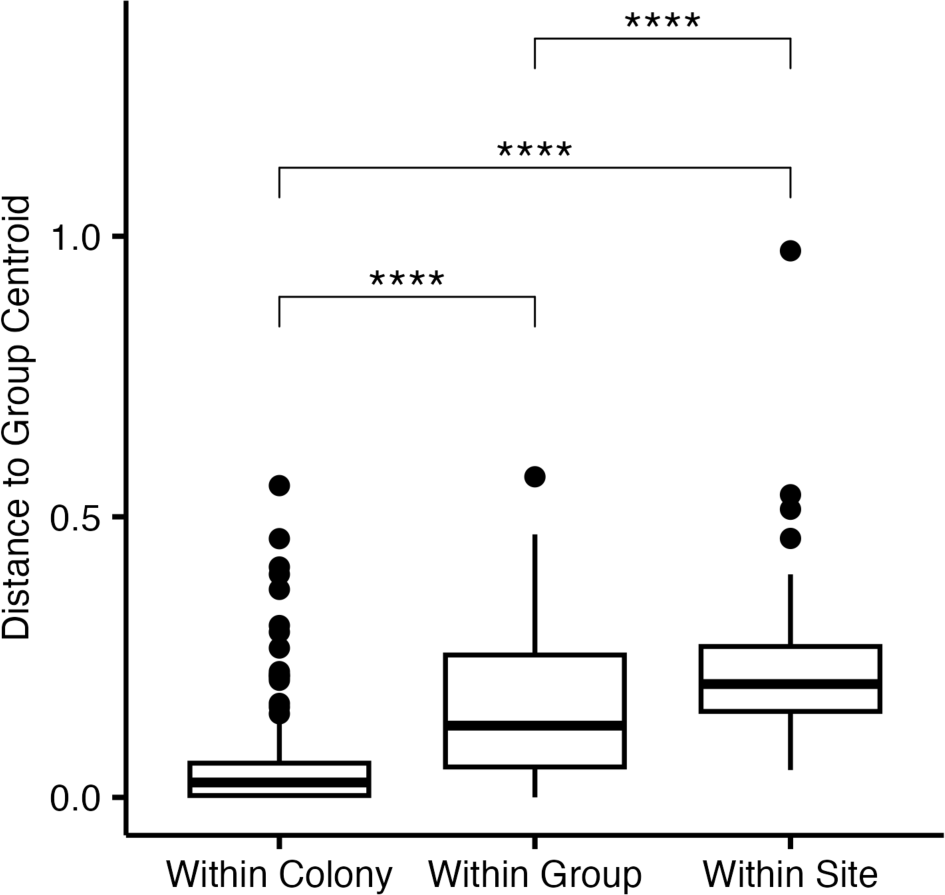
Symbiodiniaceae within colony variance less than within group and within site variance. Community variance determined by distances to centroid in multivariate space. Significance determined by a pairwise Wilcox test with a significance threshold of *p=0.05*.

**Supplemental Figure 5.**
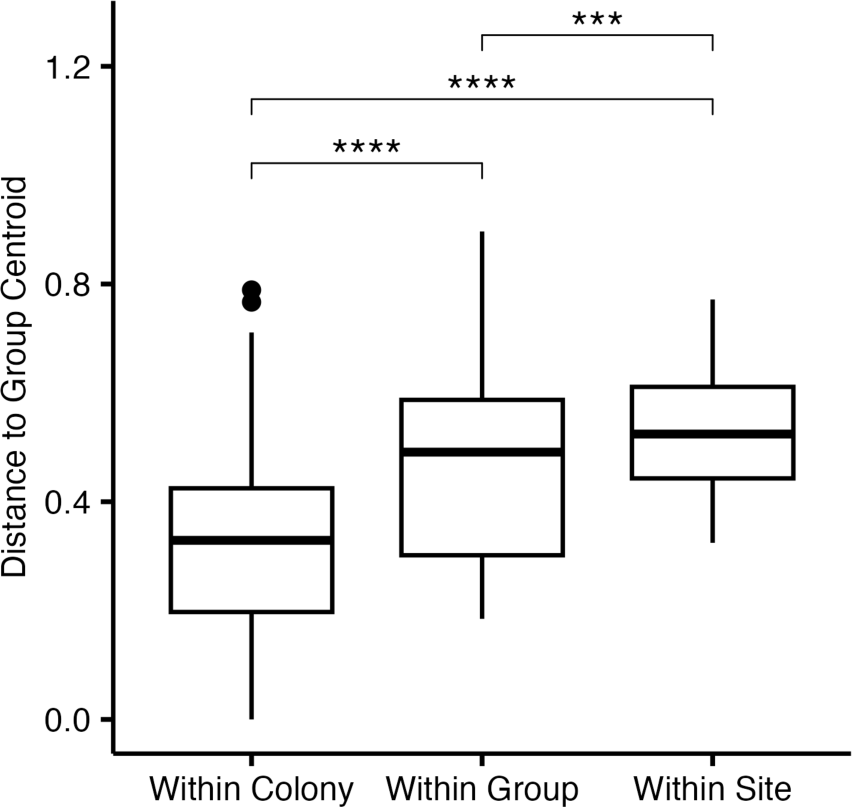
Microbial within colony variance less than within group and within site variance. Community variance determined by distances to centroid in multivariate space. Significance determined by a pairwise Wilcox test with a significance threshold of *p=0.05*.

**Supplemental Figure 6.**
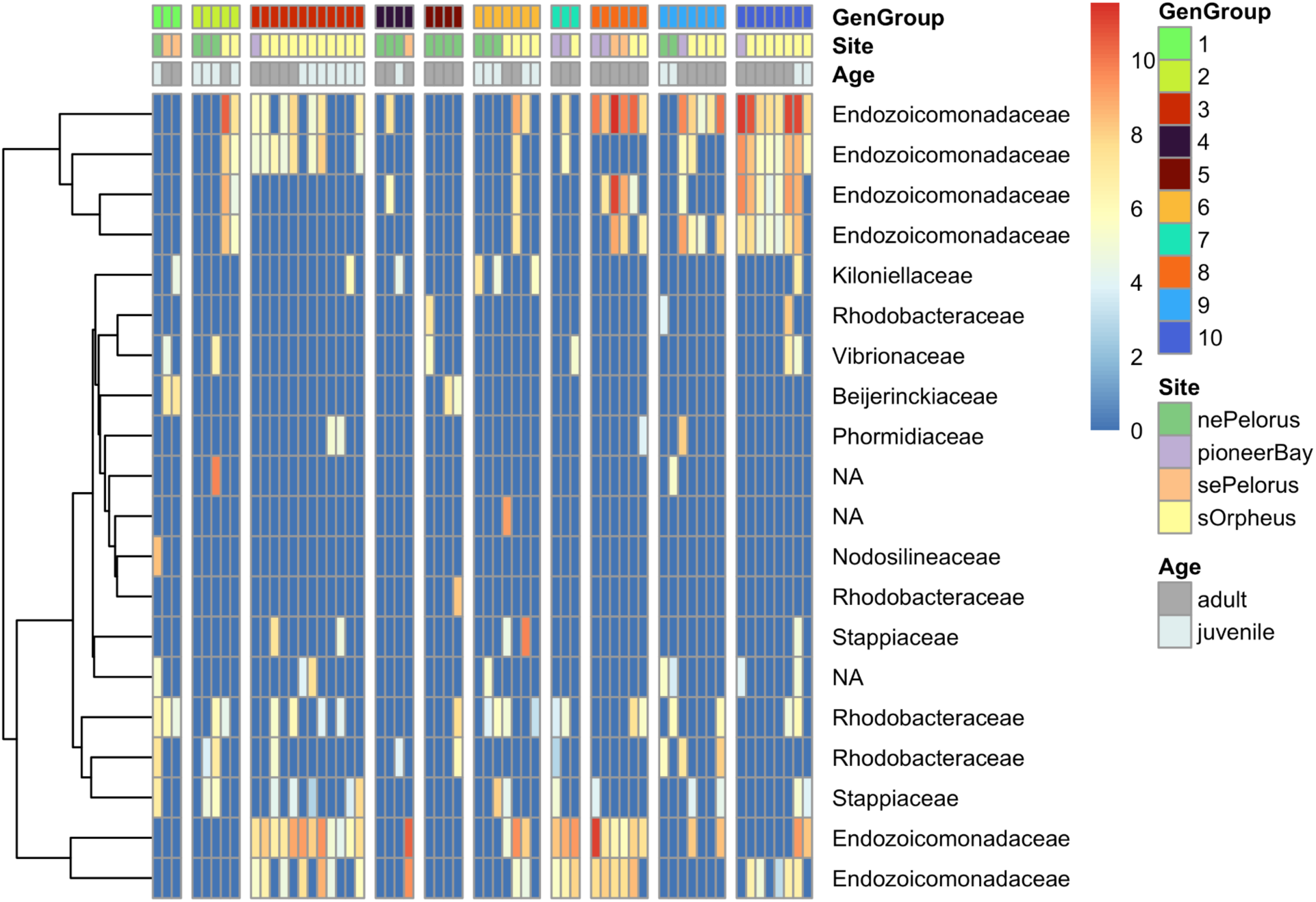
Top 20 most abundant ASVs across samples. Top average 20 most abundant bacterial ASVs occurring in samples. Fill shows abundance by log2 normalized counts. Rows show ASV abundance across samples, with assigned family indicated on the right. NA values indicate no assignment to the family level for a particular ASV. Columns reflect individual samples, separated by genetic group. Site and size class for each sample are indicated at the top of the plot.

**Supplemental Figure 7.**
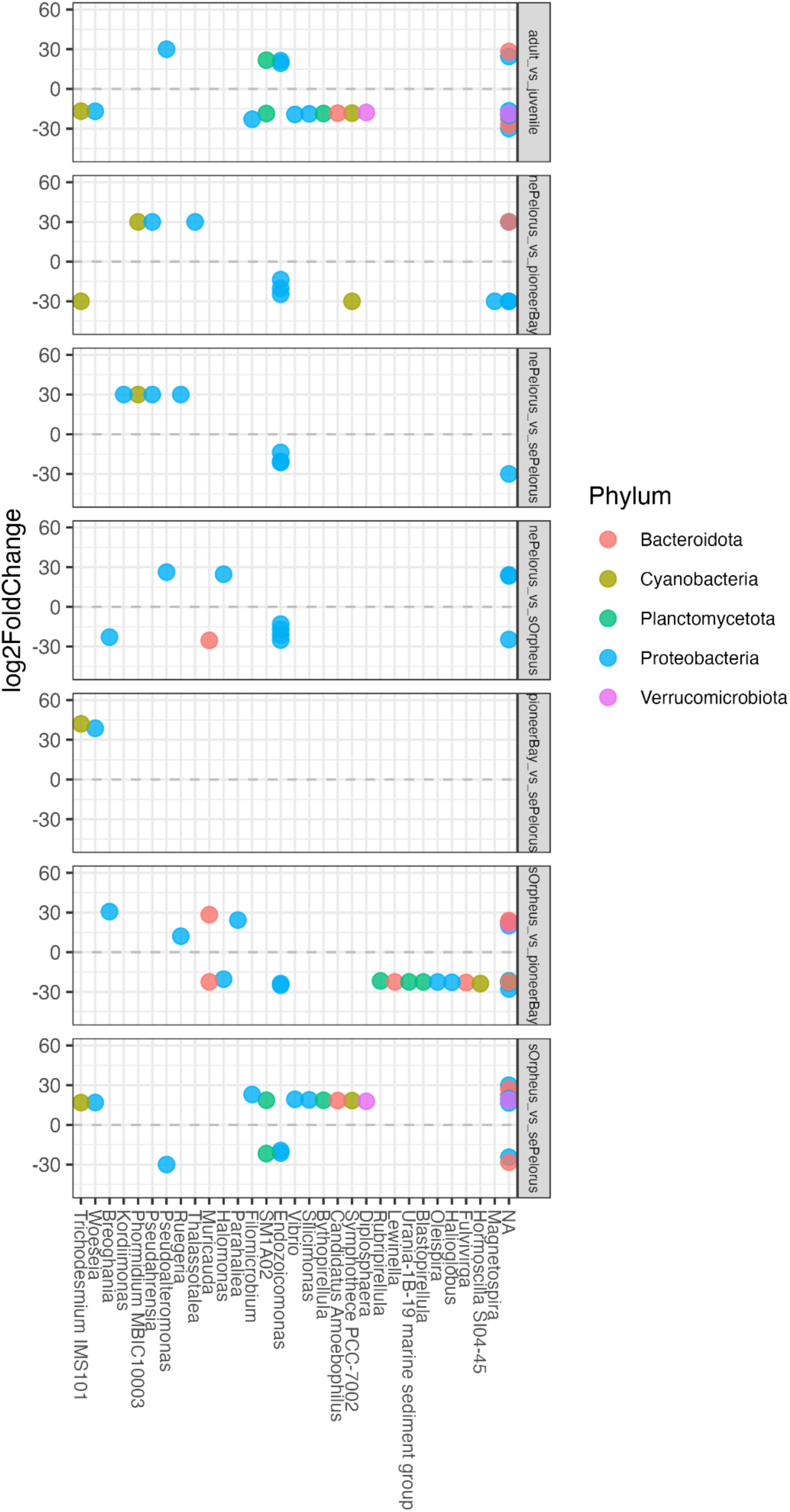
Significantly differentially abundant ASVs across all comparisons. Panels give pairwise comparisons, by age class and site, of significantly differentially abundant ASVs at the α < 0.01 level. Columns give genus-level assignments, colors indicate phylum-level assignment.

## Supplemental Tables

**Table 1.** PERMANOVA results for drivers of host genetic structure within each admixture group. The only significant predictor (at the *p < 0.01* level, when corrected for multiple hypothesis testing) is site for admixture group 4.

|  | Admix_1 |  | Admix_2 |  | Admix_3 |  | Admix_4 |  | Admix_5 |  |
| --- | --- | --- | --- | --- | --- | --- | --- | --- | --- | --- |
| Variable | $R^2$ | $p$ | $R^2$ | $p$ | $R^2$ | $p$ | $R^2$ | $p$ | $R^2$ | $p$ |
| Site | 0.087 | 0.392 | 0.320 | 0.818 | 0.084 | 0.087 | **0.100** | **0.005\*** | 0.333 | 0.583 |
| Size | 0.047 | 0.213 | NA | NA | 0.103 | 0.050 | 0.040 | 0.490 | 0.336 | 0.500 |
| Residual | .719 |  | 0.679 |  | 0.492 |  | 0.860 |  | 0.331 |  |

**Table 2.** Symbiodiniaceae community gradient forest model importances with genetic subcluster as a predictor. Overall gradient forest model importance (proportion variance explained) and the importance of each predictor. Higher importance corresponds to better predictive power, and an overall better model.

| Predictor | Importance |
| --- | --- |
| Subcluster | 0.155 |
| Site | 0.021 |
| Size Class | 0.016 |
| Total Importance | 0.193 |

**Table 3.** Microbial community gradient forest model importances with genetic subcluster as a predictor. Overall gradient forest model importance (proportion variance explained) and the importance of each predictor. Higher importance corresponds to better predictive power, and an overall better model.

| Predictor | Importance |
| --- | --- |
| Subcluster | 0.212 |
| Site | 0.148 |
| Size Class | 0.050 |
| Total | 0.425 |
| Importance |  |

**Table 4.** Symbiodiniaceae community gradient forest model importances with admixture group as a predictor. Overall gradient forest model importance (proportion variance explained) and the importance of each predictor. Higher importance corresponds to better predictive power, and an overall better model. Results are given in the main text for the model including genetic subcluster as a predictor (Supp Table 2) as it has higher overall importance.

| Predictor | Importance |
| --- | --- |
| AdmixGroup | 0.100 |
| Site | 0.023 |
| Size Class | 0.011 |
| Total Importance | 0.135 |

**Table 5.**
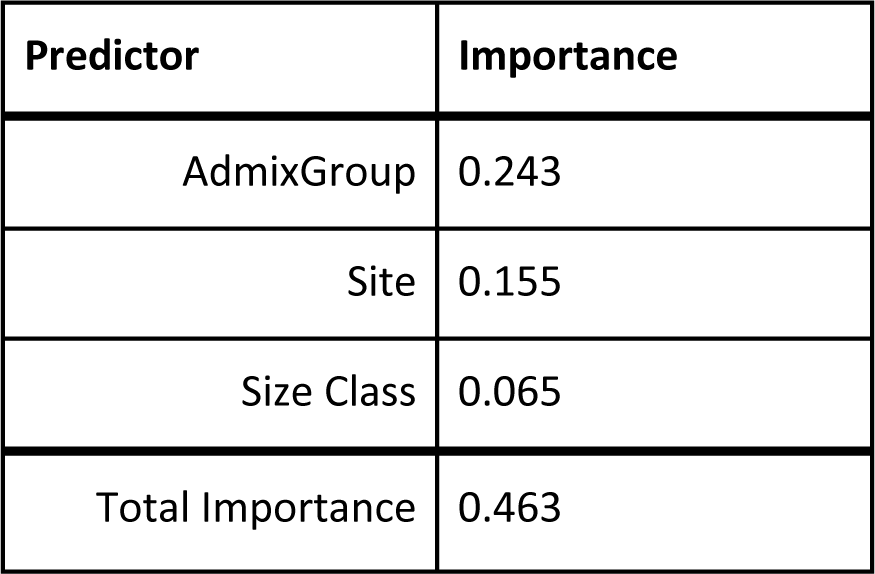
Microbial community gradient forest model importances with admixture group as a predictor. Overall gradient forest model importance (proportion variance explained) and the importance of each predictor. Higher importance corresponds to better predictive power, and an overall better model. Results are given in the main text for the model including genetic subcluster as a predictor (Supp Table 3) for consistency with the Symbiodiniaceae model.

**Table 6.** PERMANOVA results for drivers of Symbiodiniaceae community structure using admixture group rather than genetic subcluster. Admixture group still has the strongest (and only significant) effect on symbiont community.

| Predictor | $R^2$ | $F$ | $p$ |
| --- | --- | --- | --- |
| AdmixGroup | 0.394 | 7.42 | 0.001 |
| Site | 0.059 | 1.48 | 0.146 |
| Size Class | 0.015 | 1.16 | 0.288 |
| Residual | 0.531 |  |  |

**Table 7.**
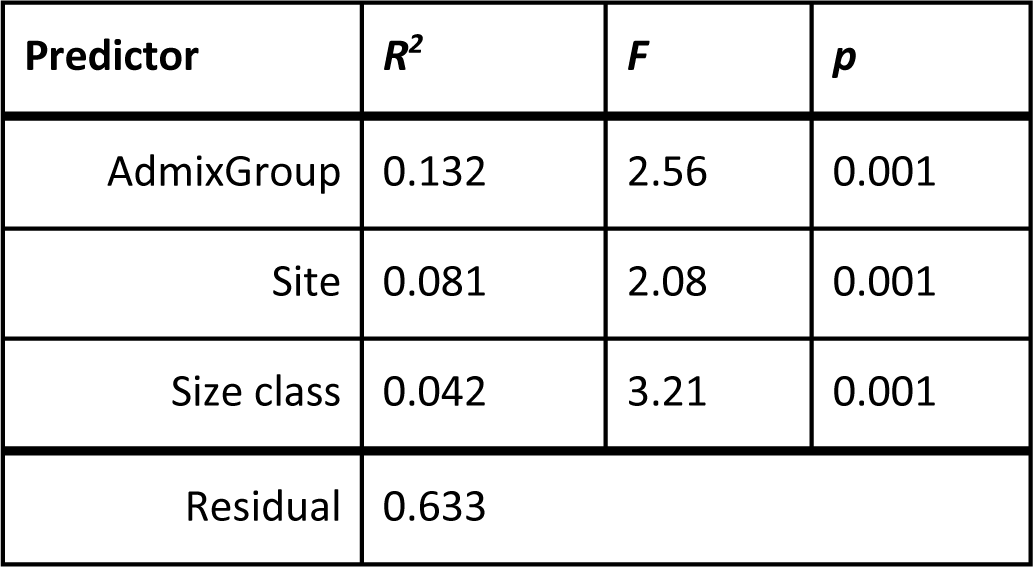
PERMANOVA results for drivers of microbial community structure using admixture group rather than genetic subcluster. admixture group still has the strongest effect on symbiont community, followed by a significant effect of site and size class.

## References

Baker, A. C. (2003). Flexibility and Specificity in Coral-Algal Symbiosis: Diversity, Ecology, and Biogeography of Symbiodinium. *Annual Review of Ecology*, Evolution, and Systematics, 34(1), 661–689. 10.1146/annurev.ecolsys.34.011802.132417

Bengtsson-Palme, J., Hartmann, M., Eriksson, K. M., Pal, C., Thorell, K., Larsson, D. G. J., & Nilsson, R. H. (2015). metaxa2: Improved identification and taxonomic classification of small and large subunit rRNA in metagenomic data. Molecular Ecology Resources, 15(6), 1403–1414. 10.1111/1755-0998.12399

Ben-Haim, Y., Zicherman-Keren, M., & Rosenberg, E. (2003). Temperature-Regulated Bleaching and Lysis of the Coral Pocillopora damicornis by the Novel Pathogen Vibrio coralliilyticus. Applied and Environmental Microbiology, 69(7), 4236–4242. 10.1128/AEM.69.7.4236-4242.2003

Berkelmans, R., & van Oppen, M. J. H. (2006). The role of zooxanthellae in the thermal tolerance of corals: A ‘nugget of hope’ for coral reefs in an era of climate change. Proceedings of the Royal Society B: Biological Sciences, 273(1599), 2305–2312. 10.1098/rspb.2006.3567

Black, K., Rippe, J., & Matz, M. (2022). *Environmental drivers of genetic adaptation in Florida corals* [Preprint]. Preprints. 10.22541/au.166997531.17274551/v1

Brown, D. P., Basch, L., Barshis, D., Forsman, Z., Fenner, D., & Goldberg, J. (2009). American Samoa’s island of giants: Massive Porites colonies at Ta’u island. Coral Reefs, 28(3), 735–735. 10.1007/s00338-009-0494-8

Callahan, B. J., McMurdie, P. J., Rosen, M. J., Han, A. W., Johnson, A. J. A., Holmes, S. P. (2016). DADA2: High resolution sample inference from Illumina amplicon data. Nature Methods, 13(7), 581–583. 10.1038/nmeth.3869

Claar, D. C., Starko, S., Tietjen, K. L., Epstein, H. E., Cunning, R., Cobb, K. M., Baker, A. C., Gates, R. D., & Baum, J. K. (2020). Dynamic symbioses reveal pathways to coral survival through prolonged heatwaves. Nature Communications, 11(1), 6097. 10.1038/s41467-020-19169-y

Claar, D. C., Tietjen, K. L., Cox, K. D., Gates, R. D., & Baum, J. K. (2020). Chronic disturbance modulates symbiont (Symbiodiniaceae) beta diversity on a coral reef. Scientific Reports, 10(1), 4492. 10.1038/s41598-020-60929-z

Coffroth, M. A., Leigh, N. J., McIlroy, S. E., Miller, M. W., & Sheets, H. D. (2022). Genetic structure of dinoflagellate symbionts in coral recruits differs from that of parental or local adults. Ecology and Evolution, 12(9), e9312. 10.1002/ece3.9312

Dougan, K. E., Bellantuono, A. J., Kahlke, T., Abbriano, R. M., Chen, Y., Shah, S., Granados-Cifuentes, C., van Oppen, M. J. H., Bhattacharya, D., Suggett, D. J., Chan, C. X., & Rodriguez-Lanetty, M. (2022). Whole-genome duplication in an algal symbiont serendipitously confers thermal tolerance to corals [Preprint]. Genomics. 10.1101/2022.04.10.487810

Dunphy, C. M., Gouhier, T. C., Chu, N. D., & Vollmer, S. V. (2019). Structure and stability of the coral microbiome in space and time. Scientific Reports, 9(1), 1–13. 10.1038/s41598-019-43268-6

Ellis, N., Smith, S. J., & Pitcher, C. R. (2012). Gradient forests: Calculating importance gradients on physical predictors. Ecology, 93(1), 156–168. 10.1890/11-0252.1

Epstein, H. E., Smith, H. A., Cantin, N. E., Mocellin, V. J. L., Torda, G., & van Oppen, M. J. H. (2019). Temporal Variation in the Microbiome of Acropora Coral Species Does Not Reflect Seasonality. Frontiers in Microbiology, 10. https://www.frontiersin.org/article/10.3389/fmicb.2019.01775

Fifer, J. E., Yasuda, N., Yamakita, T., Bove, C. B., & Davies, S. W. (2022). Genetic divergence and range expansion in a western North Pacific coral. Science of The Total Environment, 813, 152423. 10.1016/j.scitotenv.2021.152423

Forsman, Z. H., Ritson-Williams, R., Tisthammer, K. H., Knapp, I. S. S., & Toonen, R. J. (2020). Host-symbiont coevolution, cryptic structure, and bleaching susceptibility, in a coral species complex (Scleractinia; Poritidae). Scientific Reports, 10(1), Article 1. 10.1038/s41598-020-73501-6

Glasl, B., Bourne, D. G., Frade, P. R., Thomas, T., Schaffelke, B., & Webster, N. S. (2019). Microbial indicators of environmental perturbations in coral reef ecosystems. Microbiome, 7(1), 1–13. 10.1186/s40168-019-0705-7

Goulet, T. L., Erill, I., Ascunce, M. S., Finley, S. J., & Javan, G. T. (2020). Conceptualization of the Holobiont Paradigm as It Pertains to Corals. Frontiers in Physiology, 11, 566968. 10.3389/fphys.2020.566968

Grupstra, C. G. B., Meyer-Kaiser, K. S., Bennet, M. J., Andes, M. O., Juszkiewicz, D. J., Fifer, J. E., De-Anoy, J. P., Gomez-Campo, K., Martinez-Rugerio, I., Aichelman, H. E., Huzar, A. E., Hughes, A. M., Rivera, H. E., Davies, S. W. (2024). Distinct modes of holobiont specialization among cryptic coral lineages. bioRxiv. 10.1101/2024.07.05.598868

Hernandez-Agreda, A., Leggat, W., Bongaerts, P., Herrera, C., & Ainsworth, T. D. (2018). Rethinking the Coral Microbiome: Simplicity Exists within a Diverse Microbial Biosphere. mBio, 9(5), 10.1128/mbio.00812-18. https://doi.org/10.1128/mbio.00812-18

Hoadley, K. D., Pettay, Daniel. T., Lewis, A., Wham, D., Grasso, C., Smith, R., Kemp, D. W., LaJeunesse, T., & Warner, M. E. (2021). Different functional traits among closely related algal symbionts dictate stress endurance for vital Indo-Pacific reef-building corals. Global Change Biology, 27(20), 5295– 5309. 10.1111/gcb.15799

Hoegh-Guldberg, O., Mumby, P. J., Hooten, A. J., Steneck, R. S., Greenfield, P., Gomez, E., Harvell, C. D., Sale, P. F., Edwards, A. J., Caldeira, K., Knowlton, N., Eakin, C. M., Iglesias-Prieto, R., Muthiga, N., Bradbury, R. H., Dubi, A., & Hatziolos, M. E. (2007). Coral reefs under rapid climate change and ocean acidification. Science (New York, N.Y.), 318(5857), 1737–1742. 10.1126/science.1152509

Hume, B. C. C., D’Angelo, C., Smith, E. G., Stevens, J. R., Burt, J., & Wiedenmann, J. (2015). Symbiodinium thermophilum sp. Nov., a thermotolerant symbiotic alga prevalent in corals of the world’s hottest sea, the Persian/Arabian Gulf. Scientific Reports, 5(1), Article 1. 10.1038/srep08562

Hume, B. C. C., Smith, E.G., Ziegler, M., et al. (2019). SymPortal: A novel analytical framework and platform for coral algal symbiont next-generation sequencing *ITS2* profiling. Molecular Ecology Resources, 19(4), 1063–1080. 10.1111/1755-0998.13004

Korneliussen, T. S., Albrechtsen, A., & Nielsen, R. (2014). ANGSD: Analysis of Next Generation Sequencing Data. BMC Bioinformatics, 15(1), 356. 10.1186/s12859-014-0356-4

Kuhn, M. (2022). *caret: Classification and Regression Training*. (6.0-92) [R package].

Ladner, J. T., & Palumbi, S. R. (2012). Extensive sympatry, cryptic diversity and introgression throughout the geographic distribution of two coral species complexes. Molecular Ecology, 21(9), 2224– 2238. 10.1111/j.1365-294X.2012.05528.x

LaJeunesse, T. C., Smith, R. T., Finney, J., & Oxenford, H. (2009). Outbreak and persistence of opportunistic symbiotic dinoflagellates during the 2005 Caribbean mass coral ‘bleaching’ event. Proceedings of the Royal Society B: Biological Sciences, 276(1676), 4139–4148. 10.1098/rspb.2009.1405

Langmead, B., & Salzberg, S. L. (2012). Fast gapped-read alignment with Bowtie 2. Nature Methods, 9(4), Article 4. 10.1038/nmeth.1923

Liaw, A., & Wiener, M. (2002). Classification and Regression by randomForest. R News, 2(3), 18–22.

Love, M. I., Huber, W., Anders, S. (2014). Moderated estimation of fold change and dispersion for RNA-seq data with DESeq2. Genome Biology, 15(12), 550.

Martin, M. (2011). Cutadapt removes adapter sequences from high-throughput sequencing reads. EMBnet.Journal, 17(1), Article 1. 10.14806/ej.17.1.200

McDevitt-Irwin, J. M., Garren, M., McMinds, R., Vega Thurber, R., & Baum, J. K. (2019). Variable interaction outcomes of local disturbance and El Niño-induced heat stress on coral microbiome alpha and beta diversity. Coral Reefs, 38(2), 331–345. 10.1007/s00338-019-01779-8

McIlroy S.E., Cunning R., Baker A.C., Coffroth M.A. (2019). Competition and succession among coral endosymbionts. Ecology and Evolution, 9, 12767–12778. 10.1002/ece3.5749

McMurdie, P. J., & Holmes, S. (2014). Waste Not, Want Not: Why Rarefying Microbiome Data Is Inadmissible. PLOS Computational Biology, 10(4), e1003531. 10.1371/journal.pcbi.1003531

Meisner, J., & Albrechtsen, A. (2018). Inferring population structure and admixture proportions in low-depth NGS data. Genetics, 210(2), 719–731. 10.1534/genetics.118.301336

Morikawa, M. K., & Palumbi, S. R. (2019). Using naturally occurring climate resilient corals to construct bleaching-resistant nurseries. Proceedings of the National Academy of Sciences, 116(21), 10586– 10591. 10.1073/pnas.1721415116

Morrow, K., Moss, A., Chadwick, N., & Liles, M. (2012). Bacterial Associates of Two Caribbean Coral Species Reveal Species-Specific Distribution and Geographic Variability. Applied and Environmental Micribiology, 78(18), 6438–6449.

Morrow, K., Pankey, M. S., Lesser, M. P., (2022). Community structure of coral microbiomes is dependent on colony morphology. Microbiome, 10(113). 10.1186/s40168-022-01308-w

Munn, C. B. (2015). The Role of Vibrios in Diseases of Corals. Microbiology Spectrum, 3(4), 10.1128/microbiolspec.ve-0006-2014. https://doi.org/10.1128/microbiolspec.ve-0006-2014

Oskansen, J., Gavin, S., & Blanchet, G. (2022). *vegan: Community Ecology Package* (2.6-2) [R package].

Peixoto, R. S., Rosado, P. M., Leite, D. C. de A., Rosado, A. S., & Bourne, D. G. (2017). Beneficial Microorganisms for Corals (BMC): Proposed Mechanisms for Coral Health and Resilience. Frontiers in Microbiology, 8. https://www.frontiersin.org/articles/10.3389/fmicb.2017.00341

Pollock, F. J., McMinds, R., Smith, S., Bourne, D. G., Willis, B. L., Medina, M., Thurber, R. V., & Zaneveld, J. R. (2018). Coral-associated bacteria demonstrate phylosymbiosis and cophylogeny. Nature Communications, 9(1), Article 1. 10.1038/s41467-018-07275-x

Quek, Z. B. R., Tanzil, J. T. I., Jain, S. S., Yong, W. L. O., Yu, D. C. Y., Soh, Z., Ow, Y. X., Tun, K., Huang, D., & Wainwright, B. J. (2023). Limited influence of seasonality on coral microbiomes and endosymbionts in an equatorial reef. Ecological Indicators, 146, 109878. 10.1016/j.ecolind.2023.109878

Rippe, J. P., Dixon, G., Fuller, Z. L., Liao, Y., & Matz, M. (2021). Environmental specialization and cryptic genetic divergence in two massive coral species from the Florida Keys Reef Tract. Molecular Ecology, 30(14), 3468–3484. 10.1111/mec.15931

Robbins, S. J. (2019). A genomic view of the reef-building coral Porites lutea and its microbial symbionts. Nature Microbiology, 4, 14.

Rouzé, H., Lecellier, G., Pochon, X., Torda, G., & Berteaux-Lecellier, V. (2019). Unique quantitative Symbiodiniaceae signature of coral colonies revealed through spatio-temporal survey in Moorea. Scientific Reports, 9(1), 7921. 10.1038/s41598-019-44017-5

Scrucca, L., Fop, M., Murphy, T. B., & Raftery, A. E. (2016). mclust 5: Clustering, Classification and Density Estimation Using Gaussian Finite Mixture Models. The R Journal, 8(1), 289–317.

Shoguchi, E., Beedessee, G., Tada, I., Hisata, K., Kawashima, T., Takeuchi, T., Arakaki, N., Fujie, M., Koyanagi, R., Roy, M. C., Kawachi, M., Hidaka, M., Satoh, N., & Shinzato, C. (2018). Two divergent Symbiodinium genomes reveal conservation of a gene cluster for sunscreen biosynthesis and recently lost genes. BMC Genomics, 19(1), 458. 10.1186/s12864-018-4857-9

Shoguchi, E., Shinzato, C., Kawashima, T., Gyoja, F., Mungpakdee, S., Koyanagi, R., Takeuchi, T., Hisata, K., Tanaka, M., Fujiwara, M., Hamada, M., Seidi, A., Fujie, M., Usami, T., Goto, H., Yamasaki, S., Arakaki, N., Suzuki, Y., Sugano, S., … Satoh, N. (2013). Draft assembly of the Symbiodinium minutum nuclear genome reveals dinoflagellate gene structure. Current Biology: CB, 23(15), 1399–1408. 10.1016/j.cub.2013.05.062

Smith, A., Cook, N., Cook, K., Brown, R., Woodgett, R., Veron, J., & Saylor, V. (2021). Field measurements of a massive Porites coral at Goolboodi (Orpheus Island), Great Barrier Reef. Scientific Reports, 11(1), 15334. 10.1038/s41598-021-94818-w

Starko, S., Fifer, J. E., Claar, D. C., Davies, S. W., Cunning, R., Baker, A. C., & Baum, J. K. (2023). Marine heatwaves threaten cryptic coral diversity and erode associations among coevolving partners. Science Advances, 9(32), eadf0954. 10.1126/sciadv.adf0954

Stat, M., Loh, W. K. W., LaJeunesse, T. C., Hoegh-Guldberg, O., & Carter, D. A. (2009). Stability of coral–endosymbiont associations during and after a thermal stress event in the southern Great Barrier Reef. Coral Reefs, 28(3), 709–713. 10.1007/s00338-009-0509-5

Tandon, K., Chiou, Y.-J., Yu, S.-P., Hsieh, H. J., Lu, C.-Y., Hsu, M.-T., Chiang, P.-W., Chen, H.-J., Wada, N., & Tang, S.-L. (2022). Microbiome Restructuring: Dominant Coral Bacterium Endozoicomonas Species Respond Differentially to Environmental Changes. mSystems, 7(4), e00359–22. 10.1128/msystems.00359-22

Thornhill, D. J., LaJeunesse, T. C., Kemp, D. W., Fitt, W. K., & Schmidt, G. W. (2006). Multi-year, seasonal genotypic surveys of coral-algal symbioses reveal prevalent stability or post-bleaching reversion. Marine Biology, 148(4), 711–722. 10.1007/s00227-005-0114-2

Voolstra, C. R., Suggett, D. J., Peixoto, R. S., Parkinson, J. E., Quigley, K. M., Silveira, C. B., Sweet, M., Muller, E. M., Barshis, D. J., Bourne, D. G., & Aranda, M. (2021). Extending the natural adaptive capacity of coral holobionts. Nature Reviews Earth & Environment, 2(11), Article 11. 10.1038/s43017-021-00214-3

Wainwright, B. J., Zhan, G. L., Afiq-Rosli, L., Tanzil, J. T. I., Huang, D. (2020). Host age is not a consistent predictor of microbial diversity in the coral *Porites lutea*. Scientific Reports, 10(14376). 10.1038/s41598-020-71117-4

Zhu, W., Liu, X., Zhu, M., Li, X., Yin, H., Huang, J., Wang, A., & Li, X. (2022). Responses of Symbiodiniaceae Shuffling and Microbial Community Assembly in Thermally Stressed Acropora hyacinthus. Frontiers in Microbiology, 13. https://www.frontiersin.org/articles/10.3389/fmicb.2022.832081

